# Optimal gap-affine alignment in *O*(*s*) space

**DOI:** 10.1101/2022.04.14.488380

**Authors:** Santiago Marco-Sola, Jordan M. Eizenga, Andrea Guarracino, Benedict Paten, Erik Garrison, Miquel Moreto

**Affiliations:** Computer Sciences Department, Barcelona Supercomputing Center, Barcelona, 08034, Spain; Departament d’Arquitectura de Computadors i Sistemes Operatius, Universitat Autònoma de Barcelona, Barcelona, 08193, Spain; Genomics Institute, University of California Santa Cruz, Santa Cruz, CA 95064, USA; Genomics Research Centre, Human Technopole, Viale Rita Levi-Montalcini 1, Milan, 20157, Italy; Department of Genetics, Genomics and Informatics, University of Tennessee Health Science Center, Memphis, TN 38163, USA; Departament d’Arquitectura de Computadors, Universitat Politècnica de Catalunya, Barcelona, 08034, Spain

## Abstract

**Motivation:** Pairwise sequence alignment remains a fundamental problem in computational biology and bioinformatics. Recent advances in genomics and sequencing technologies demand faster and scalable algorithms that can cope with the ever-increasing sequence lengths. Classical pairwise alignment algorithms based on dynamic programming are strongly limited by quadratic requirements in time and memory. The recently proposed wavefront alignment algorithm (WFA) introduced an efficient algorithm to perform exact gap-affine alignment in *O*(*ns*) time, where *s* is the optimal score and *n* is the sequence length. Notwithstanding these bounds, WFA’s *O*(*s*^2^) memory requirements become computationally impractical for genome-scale alignments, leading to a need for further improvement.

**Results:** In this paper, we present the bidirectional WFA algorithm (BiWFA), the first gap-affine algorithm capable of computing optimal alignments in *O*(*s*) memory while retaining WFA’s time complexity of *O*(*ns*). As a result, this work improves the lowest known memory bound *O*(*n*) to compute gap-affine alignments. In practice, our implementation never requires more than a few hundred MBs aligning noisy Oxford Nanopore Technologies reads up to 1 Mbp long while maintaining competitive execution times.

**Availability:** All code is publicly available at https://github.com/smarco/BiWFA-paper

## 1 Introduction

Pairwise sequence alignment provides a parsimonious transformation of one string into another. From this transformation, we can understand the relationship between pairs of sequences. Because similarities and differences between biosequences (DNA, RNA, protein) relate to variation in function and evolutionary history of living things, pairwise sequence alignment algorithms are a core part of many essential bioinformatics methods in read mapping (Li, 2013; Marco-Sola *et al*., 2012), genome assembly (Simpson *et al*., 2009; Koren *et al*., 2017), variant calling (Garrison and Marth, 2012; McKenna *et al.*, 2010; Rodríguez-Martín et al., 2017), and many others (Durbin et al., 1998; Jones et al., 2004). Its importance has motivated the research and development of multiple solutions over the past 50 years.

Classical approaches to derive alignments involve the application of *dynamic programming* (DP) techniques. These methods require computing a matrix whose dimensions correspond to the lengths of the query *q* and target *t* sequences. Using DP recurrence relations, these methods compute the optimal alignment score for progressively longer prefixes of q and t, which correspond to the cells of the DP matrix. Thus, an optimal alignment can then be read out by tracing the recurrence back through the matrix.

Selecting a suitable alignment score function is essential to obtain biologically meaningful alignments, as it determines the characteristics of optimal alignments. In effect, the alignment score function encodes prior expectations about the probability of certain kinds of sequence differences. It has been observed that, in many contexts, insertions and deletions are non-uniformly distributed; they are infrequent but tend to be adjacent so that they form extended *gaps* with a long-tailed length distribution. This motivated the development of *gap-affine* models in which the penalty of starting a new gap is larger than that of extending a gap (Gotoh, 1982). Crucially, gap-affine penalties can be implemented efficiently using additional DP matrices.

Problematically, the efficiency of classical gap-affine DP-based methods is constrained by their quadratic requirements in time and memory with respect to the lengths of the sequence pair. Consequently, multiple optimizations have been proposed over the years. Notable examples include bit-parallel techniques (Loving *et al*., 2014), data-layout transformations to exploit SIMD instructions (Rognes and Seeberg, 2000; Farrar, 2007; Wozniak, 1997), difference encoding of the DP matrix (Suzuki and Kasahara, 2018), among other methods (Altschul *et al*., 1990; Kiełbasa *et al*., 2011; Xia *et al*., 2021; Zhao *et al*., 2013). Nonetheless, all these exact methods retain the quadratic requirements of the original DP algorithm and therefore struggle to scale when aligning long sequences.

In many cases, when two sequences are homologous, the majority of possible alignments are largely sub-optimal, having a substantially worse score than the optimal one. For this reason, heuristic methods are usually employed to find candidate alignment regions when the cost of exact algorithms becomes impractical. Most notable approaches use adaptive *band* methods (Suzuki and Kasahara, 2017) or pruning strategies (e.g., X-drop (Zhang *et al*., 2000) and Z-drop (Li, 2018)) to avoid the computation of alignments extremely unlikely to be optimal. These heuristic methods have been implemented within many widely-used tools (Altschul *et al.*, 1990; Li, 2018).

Recently, we proposed the wavefront alignment algorithm (WFA) (Marco-Sola *et al*., 2021) to compute the exact alignment between two sequences using gap-affine penalties. WFA reformulates the alignment problem to compute the longest-possible alignments of increasing score until the optimal alignment is found. Notably, WFA takes advantage of homologous regions between sequences to accelerate alignment’s computation. As a result, WFA computes optimal gap-affine alignments in *O*(*ns*) time and *O*(*s*^2^) memory, where *n* is the sequence length and *s* the optimal alignment score. Being an exact algorithm, WFA provides the same guarantee for optimality as classical algorithms (Needleman and Wunsch, 1970; Smith and Waterman, 1981; Gotoh, 1982), but does away with the quadratic requirements in time.

WFA unlocked the path for optimal alignment methods capable of scaling to long sequences. Nevertheless, the *O*(*s*^2^) memory requirements quickly become the limiting factor when aligning sufficiently long or noisy sequences (Eizenga and Paten, 2022). As it happens, WFA’s memory requirements can be impractical when aligning through large structural variations or highly divergent genome regions. Given that we use alignment to understand variation, these are some contexts in which optimal alignment could be most useful, but its memory requirements make it prohibitive.

To address this problem, this paper presents the first gap-affine alignment algorithm to compute the optimal alignment in *O*(*ns*) time and *O*(*s*) memory (excluding the storage of the input sequences). Our method, the bidirectional WFA algorithm (BiWFA), computes the WFA alignment of two sequences in the forward and reverse direction until they meet. Using two wavefronts of *O*(*s*) memory, we demonstrate how to find the optimal breakpoint of the alignment at score ~ *s*/2 and proceed recursively to solve the complete alignment in *O*(*ns*) time. To our knowledge, this work improves the lowest known memory bound to compute gap-affine alignments *O*(*n*) (Myers and Miller, 1988) to *O*(*s*), while retaining the time complexity of the original WFA *O*(*ns*). Furthermore, our experimental results demonstrate that the BiWFA delivers comparable, or even better, performance than the original WFA, outperforming other state-of-the-art tools while using a minimal amount of memory.

The rest of the paper is structured as follows. Section 2 presents the definitions, algorithms, and formal proofs supporting BiWFA. Section 3 shows the experimental evaluation of our method, comparing it against other state-of-the-art tools and libraries. Lastly, Section 4 presents a discussion on the BiWFA method and summarizes the contributions and impact of this work.

## 2 Methods

### 2.1 Wavefront alignment algorithm

Let the query *q* = *q*_0_*q*_1_ … *q*_*n*-1_ and the text *t* = *t*_0_*t*_1_ … *t*_*m*-1_ be strings of length *n* and *m*, respectively. Likewise, let *v*[*i,j*] = *v_i_v*_*i*+1_ …*v_j_* denote a substring of any string *v* from the *i*-th to the *j*-th character. We will use (*x, o, e*) to denote the gap-affine penalties. A mismatch costs *x*, and a gap of length *l* costs *o* + *l* · *e*. We assume that *x* > 0 and *e* > 0, and further that all of the score parameters are constants.

Basically, WFA computes partial optimal alignments of increasing score until an alignment with score *s* reaches coordinate (*n, m*) of the DP matrix. In this way, the algorithm determines that *s* is the minimal alignment score. Moreover, it can derive the optimal alignment by tracing back the partial alignments that led to score *s* at (*n,m*).

Let 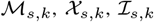, and 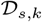 denote the offset within diagonal *k* in the DP-matrix to the farthest-reaching (f.r.) cell that has score s and ends with a match, mismatch, insertion, or deletion, respectively. In general, we denote by *wavefront* the tuple of offsets for a given score 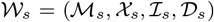. We refer to the four elements in this tuple as - its *components,* and we associate a corresponding sentinel value to specify each component: *c* ∈ {*M, X, I, D*}.

In (Marco-Sola *et al*., 2021), we proved that the f.r. points of 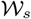 can be computed using previous wavefronts 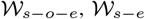, and 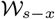, using Eq. 1 where *LCP*(*v, w*) is the length of longest common prefix between substrings *v* and *w*. The base case for this recursion is given by 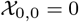.

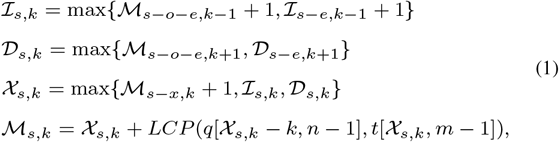

In order to compute the next wavefront, Eq. 1 shows that it is only necessary to have access to the previous *p* = max{*x, o* + *e*} wavefronts. We refer to *p* as the *scope*. Also, note that 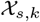 does not need to be explicitly stored as its values can be inferred using 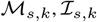, and 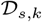.

In the worst case, WFA requires computing s wavefronts of increasing length, totalling 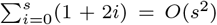 cells. Moreover, the *LCP* must be computed once for each cell. However, within a diagonal, the total number of offsets increments cannot exceed the length of the sequences. Hence, WFA requires *O*(*ns*) time and *O*(*s*^2^) memory in the worst case (Marco-Sola *et al*., 2021). Since *s* ≤ *pn*, the *O*(*ns*) factor of the execution time, due to the *LCP*, dominates over the *O*(*s*^2^) factor in the worst case. However, in practice, the time is often closer to *O*(*s*^2^ +*n*). This is because spurious matches between high-entropy sequences are short in expectation. Accordingly, the LCP computations often finishes after performing only a few character comparisons, except along the optimal alignment in which *O*(*n*) comparisons are required.

### 2.2 Bidirectional wavefront alignment algorithm

The core idea of the BiWFA algorithm is to perform WFA simultaneously in both directions on the strings: from start to end (i.e., forward) and from end to start (i.e., reverse). Each direction will only retain *p* wavefronts in memory. This is insufficient to perform a full traceback. However, when they “meet” in the middle, we can infer a breakpoint in the alignment that divides the optimal score roughly in half. Then, we can apply the same procedure on the two sides of the breakpoint recursively. We will show that this results in only a constant-factor slowdown. This technique has previously been employed to a similar end with the Myers O(ND) difference algorithm (Myers, 1986).

Figure 1 presents a graphical example of BiWFA computing a breakpoint in the optimal alignment between two sequences. The figure shows the DP cells computed by the forward and reverse wavefronts. Alignments in both directions progress until they overlap on cell (4, 4) with score 8 + 8=16 corresponding to the optimal alignment (*s_opt_* = 16).

**Fig. 1:**
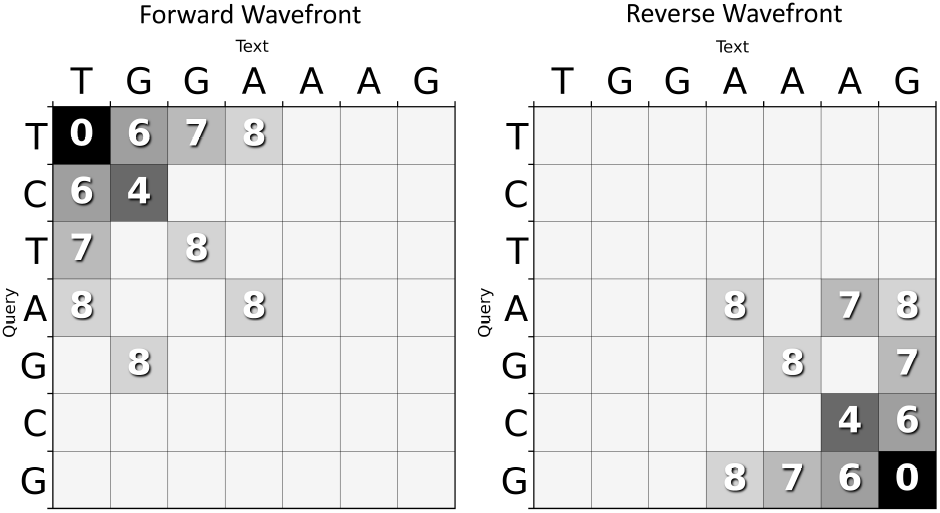
Example of BiWFA aligning *q* = “*TCTAGCG*” against *t* = “*TGGAAAG*” under the penalties (*x* = 4, *o* = 5, *e* = 1).

First, let us define the WFA equations for the forward and reverse alignment directions. The recursions for the forward direction are equivalent to those of the standard WFA presented above (Eq. 1). However, to highlight the distinction, we will denote them 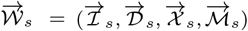. The recursions for the reverse direction are very similar (Eq. 2), using 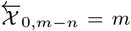 as the base case and *LCS*(*v, w*) to denote the length of the longest common suffix of *v* and *w*. Note that the same argument used in Marco-Sola *et al.* (2021) applies to the reverse recursions to prove that they are f.r. in the reverse direction.

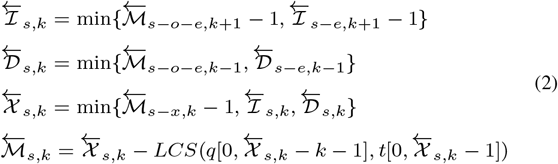

Algorithm 1 presents the BiWFA algorithm to compute a breakpoint in the optimal alignment at ~ *s*/2. Using forward and reverse wavefronts, the algorithm proceeds by alternatingly computing forward and reverse alignments (i.e., 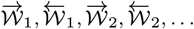). To this end, BiWFA relies on the operators *WF_NEXT()* and *WF_EXTEND()* from the standard WFA (see Marco-Sola *et al.* (2021)) to compute successive wavefronts using Eqs. 1 and 2. The process is halted after their offsets overlap to compute the position of a breakpoint in the optimal alignment. This algorithm iterates until it is guaranteed that the optimal breakpoint has been found. However, there are some technical details involving the detection of overlaps and the computation of the optimal breakpoint, which we cover in Sections 2.3 and 2.4.

**Algorithm 1:**
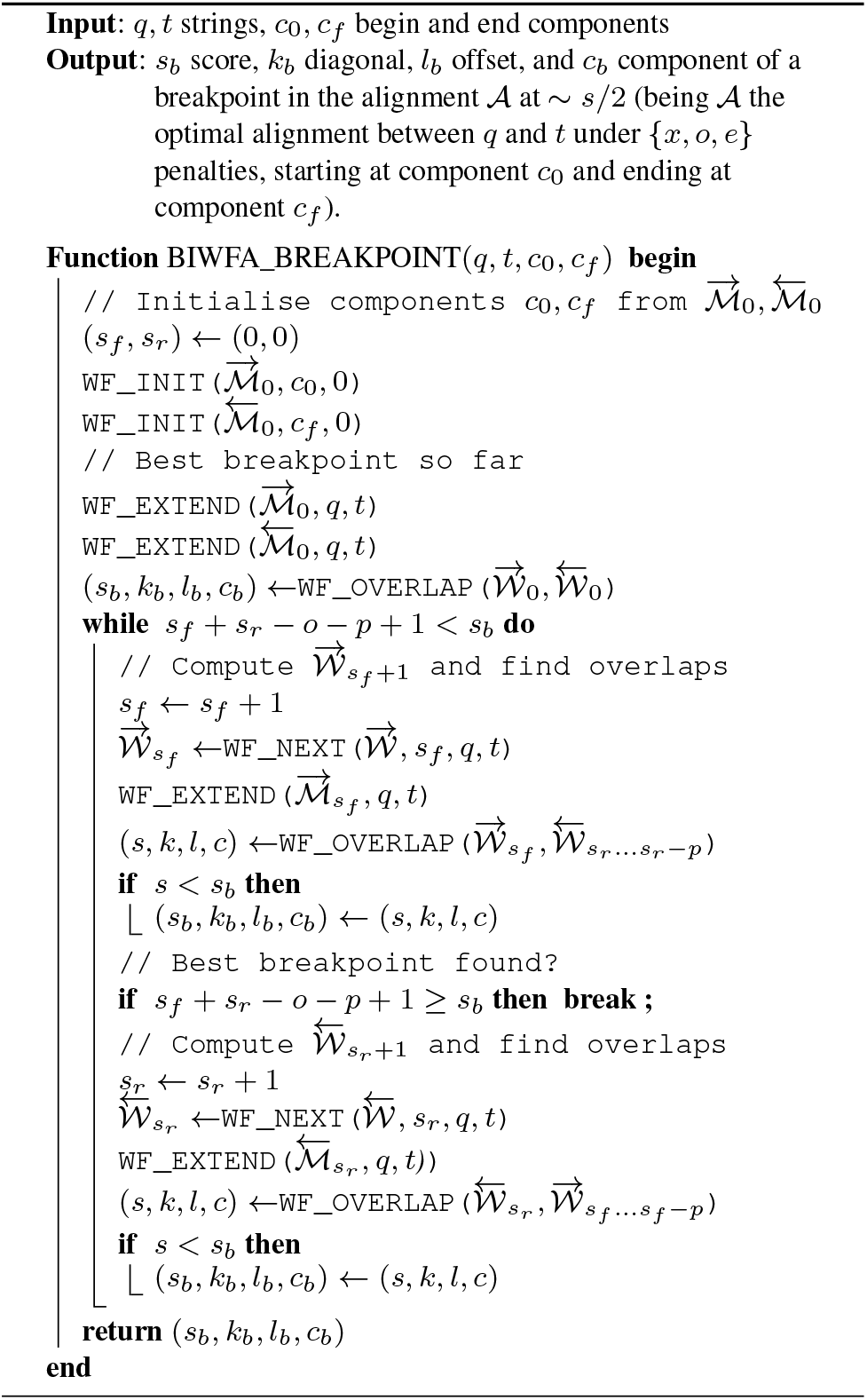
Compute optimal alignment breakpoint using BiWFA.

### 2.3 Finding a score-balanced breakpoint in the optimal alignment

The first technical detail involved in finding an alignment breakpoint between the two directions is that it is often not possible to split an alignment into an equally-scoring prefix and suffix. In general, two prefixes of the optimal alignment that differ by one character can have scores that differ by as much as *p*. Accordingly, we will demand a weaker notion of balance. If *s_f_* and *s_r_* are the forward and reverse scores, respectively, we will aim to have |*s_f_* – *s_r_*| ≤ *p*.

The second technical detail is that the optimal score is not always the sum of the two scores. This occurs because the forward iteration incurs the gap open penalty o at the beginning of gaps, but the reverse incurs it at the end of gaps (or rather, at the beginning in the reverse direction). Thus, if the two directions meet in a gap, then we have *s_opt_* = *s_f_* + *s_r_* – *o* rather than *s_opt_* = *s_f_* + *s_r_*, where *s_opt_* is the optimal alignment score.

The final technical detail is that offsets of the two directions may not precisely meet. WFA proceeds by greedily taking matches in both directions. This makes it possible for the two directions to shoot past each other without actually meeting. It turns out that it is sufficient to detect that such an overshoot has occurred, as will be shown in Section 2.4.

In Algorithm 2, we reconcile these three difficulties. Without loss of generality, we assume that a forward wavefront 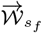 has been computed (Algorithm 1), and we want to detect overlaps against the previously computed reverse wavefronts 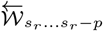. First, if 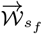 belongs to a score-balanced breakpoint (with |*s_f_* – *s_r_*| ≤ *p*), it is sufficient to check for overlaps against 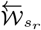 and the previous *p* – 1 reverse wavefronts. Second, for every diagonal *k* in wavefront 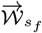, Algorithm 2 checks of overlaps in all wavefront components. This way, the algorithm keeps track of the overlap with the minimum score detected so far. Last, note that overlaps on 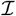 and 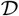 components account twice for the gap-open score *o*. Hence, the score from overlaps at indel components has to be decreased by *o*.

**Algorithm 2:**
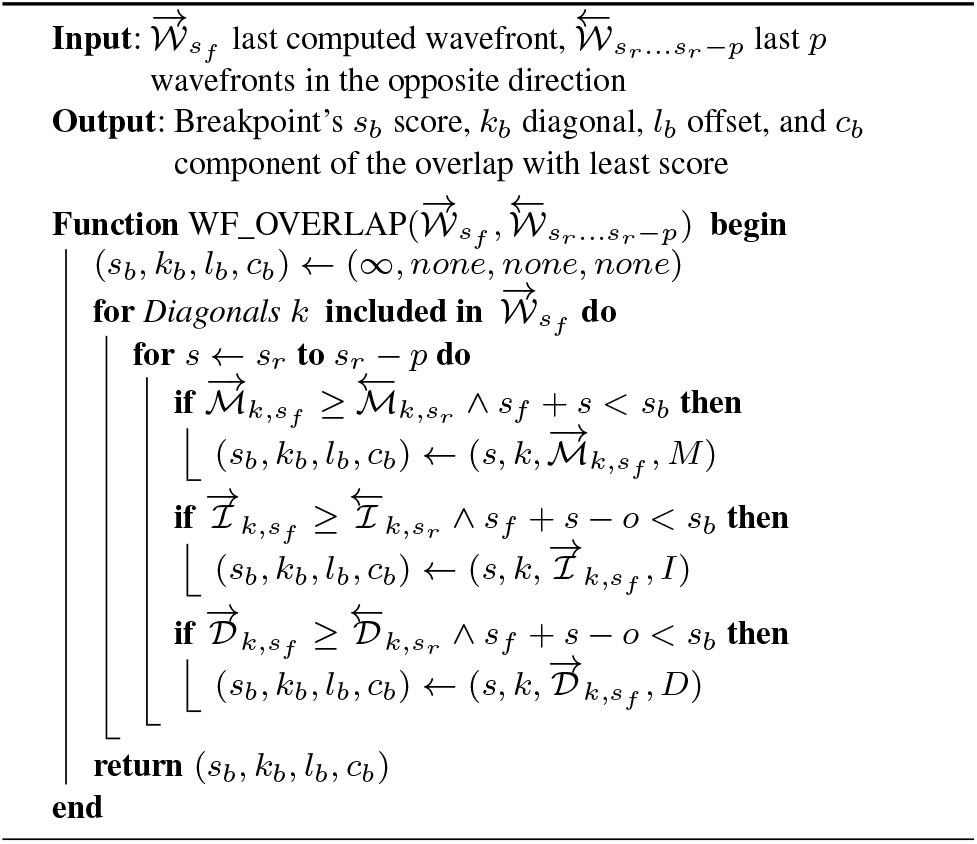
Detect overlaps and compute optimal breakpoint.

In practice, Algorithm 1 can avoid most calls to *WF_OVERLAP()*. An efficient implementation can keep track of the farthest reached antidiagonal by each wavefront. If the most advanced antidiagonal reached by the forward and reverse wavefronts do not overlap, it follows that no offsets from any diagonal can overlap, rendering the call to *WF_OVERLAP()* unnecessary.

### 2.4 Correctness of the breakpoint detection

The correctness of the Algorithm 1 stems from the following lemma.

#### Lemma 2.1.

*The optimal alignment score s_opt_* ≤ *s if and only if there exist s f, s_r_, and k such that* |*s_f_* – *s_r_*| ≤ *p and at least one of the following is true*:

1. *s_f_* + *s_r_* = *s and* 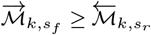
2. *s_f_* + *s_r_* = *s* + *o and* 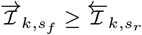
3. *s_f_* + *s_r_* = *s* + *o and* 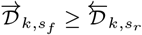,

and further, 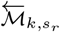 (resp. 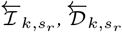) is included in the traceback of an alignment with score at most s if the first (resp. second, third) condition is true.

Proof. See supplementary material.

This lemma implies that the minimum value *s* for which the “only if” condition holds is the optimal score. Moreover, if the first of the three conditions is found to hold for some values *s_f_* and *s_r_*, then *s_opt_* ≤ *s_f_* + *s_r_* + *o*. Therefore, Algorithm 1 is guaranteed to find part of a minimum-scoring alignment based on the following features:

- Algorithm 2 checks a window of *p* score values on each iteration.
- Algorithm 1 iterates through alternatingly increasing values of *s_f_* and *s_r_*, detecting breakpoints with scores of at least *s_f_* + *s_r_* — *o* – *p* +1 in each iteration.
- After finding some *s_f_* and *s_r_* that satisfy the overlap condition, Algorithm 1 continues for additional iterations until it is no longer possible to find a lower score.

### 2.5 Combining breakpoints into an alignment

Algorithm 3 shows how to use BiWFA to recursively split alignments into smaller subproblems until the remaining alignment can be trivially solved.

**Algorithm 3:**
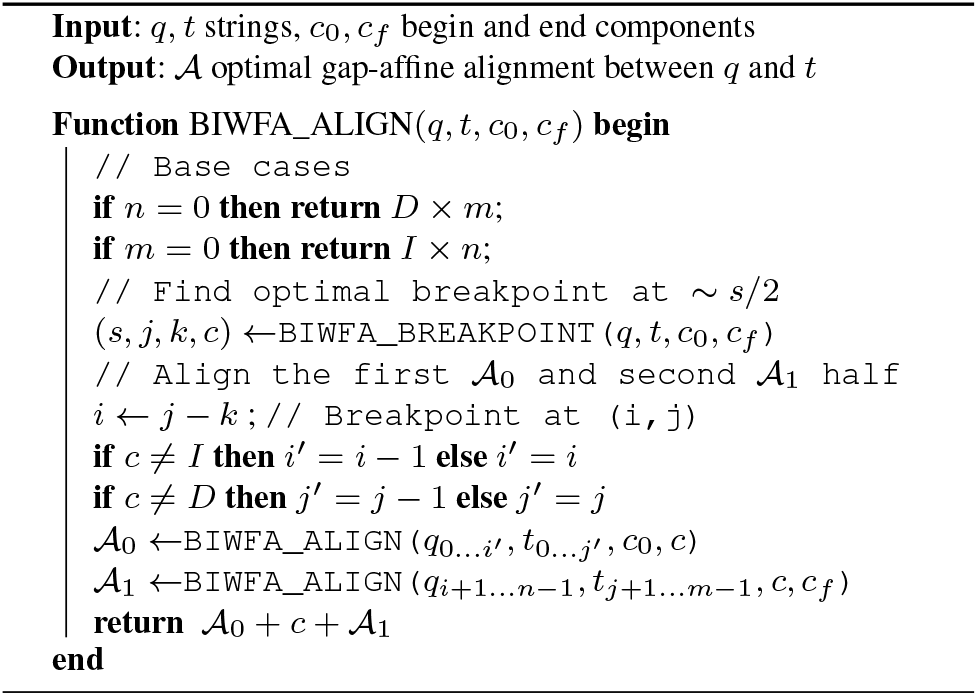
BiWFA recursive computation of the optimal alignment in *O*(*s*) space.

Note that a breakpoint computed by BiWFA can be found on the *I* or *D* components. Thus, those alignments that connect with this breakpoint have to start or end at the given component. This way, Algorithm 3 considers the starting and ending component of each alignment, and forces the underlying WFAs to use different initial condition depending on the alignment starting at the 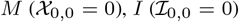, or *D* component 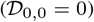. A similar argument applies to the ending conditions of each alignment ending at the 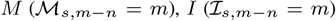, or *D* component 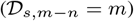.

### 2.3 BiWFA uses *O*(*s*) space and *O*((*m* + *n*)*s*) time

The memory complexity of Algorithm 1 is relatively simple to characterize. The range of diagonal values *k* increases by at most 2 every time *s* is incremented, and each forward and reverse search only needs to store the last *p* wavefronts. Thus, the memory use is proportional to the optimal alignment score, *O*(*s*), excluding the storage of the input sequences. Also, note that the output alignment only requires storing the position (*i, j*) for the mismatches, insertions, and deletions (matches can be inferred from the gaps). Concerning Algorithm 3, data structures are discarded before entering a recursive call. Therefore, the maximum memory use occurs in the outermost call, in which *s* is the optimal score of the full alignment.

The time complexity is more complicated to analyze. Our proof follows similar arguments as those from Myers (1986).

#### Theorem 2.2.

*BiWFA’s time complexity is O*((*m* + *n*)*s*), *being n and m the sequences’ length and s the optimal alignment score*.

Proof. Let *ℓ* = *m* + *n*, and let *T*(*ℓ, s*) be BiWFA’s execution time with score *s*. A call to BiWFA can result in two recursive calls. Let *ℓ_f_* and *ℓ_r_* be the combined length of the sequences in the two calls, and similarly let *s_f_* and *s_r_* be the two alignment scores. Following Lemma 2.1, we know that these variables obey the following inequalities:

- *ℓ_f_* + *ℓ_r_* ≤ *ℓ*
- *s_f_* + *s_r_* ≤ *s*
- |*s_f_* — *s_r_*| ≤ *p*

Because each direction of WFA executes in *O*(*sℓ*) time (Marco-Sola *et al*., 2021), we can choose a constant *c*_1_ large enough that the following inequality holds for all *s* > 3*p*:

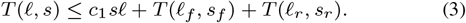

We can also choose a constant *c*_2_ large enough that for all *s* ≤ 3*p*

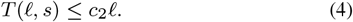

This follows because the recursion depth depends only on *s*, which we have given an upper bound. Therefore, this term includes a bounded number of calls that all have linear dependence on *ℓ*.

Next, we argue that 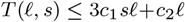 by induction on *s*. The base cases for *s* = 0,1,…, 3*p* follow trivially from the latter of the previous inequalities. Assume then that *s* > 3*p* and the induction hypothesis holds for 0,1,…, *s* – 1. Note that we then have *s_f_, s_r_* ≤ 2*s*/3, else either |*s_f_* – *s_r_*| > *p* or *s_f_* + *s_r_* > *s*. Thus,

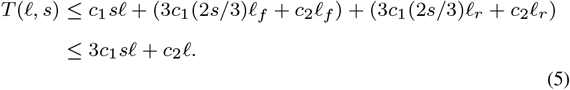

This proves the claim.

## 3 Results

We implement the BiWFA algorithm described in this work in C. The code is publicly available at https://github.com/smarco/BiWFA-paper together with the scripts required to reproduce the experimental results presented in this section.

### 3.1 Experimental setup

We evaluate the performance of our BiWFA implementation compared to the state-of-the-art and other high-performance sequence alignment libraries. We select the original WFA (Marco-Sola *et al*., 2021) (wfa-high) and its new low-memory modes (wfa-med and wfa-low) implemented in WFA2-lib (https://github.com/smarco/WFA2-lib). Also, we select the efficient wfalm (Eizenga and Paten, 2022) (wfalm) and its low-memory modes (wfalm-low and wfalm-rec). Moreover, we include the highly optimized KSW2-Z2 (ksw2_extz2_sse), from the KSW2 library (Suzuki and Kasahara, 2018; Li, 2018), as the best representative of DP-based methods due to its exceptional performance and widely usage within bioinformatics tools. Additionally, we include the Edlib (Šošić and Šikić, 2017) and BitPal (Loving *et al*., 2014) libraries, which implement bit-parallel alignment strategies for edit and non-unitary penalties (i.e., gap-linear), respectively. Although they solve a considerably easier problem (i.e., Edlib is restricted to edit-alignments and BitPal only computes the alignment score), and thus are not directly comparable, we included them in the evaluation to provide a performance upper bound.

We considered including other popular methods like those implemented in the Parasail (Daily, 2016; Wozniak, 1997; Farrar, 2007; Daily *et al*., 2015), SeqAn (Rahn *et al*., 2018), and Gaba (Suzuki and Kasahara, 2018) libraries. However, these libraries were not designed to align long and noisy sequences, and fail to complete the executions. Therefore these methods were discarded from the evaluation.

All the presented methods have been configured to generate global alignments. These algorithms are grouped in two categories: ‘Gap-affine Exact’ for exact algorithms that use gap-affine penalties (i.e., BiWFA, WFA and its low-memory modes, wfalm and its low-memory modes, and KSW2-Z2), and ‘Others’ for methods that use simpler penalty models or can only compute the alignment score (i.e., Edlib and BitPAl).

For the evaluation, we use simulated and real datasets. For the simulated datasets, we simulate several datasets of various sequence lengths (i.e., 100K, 500K, 1M, and 2M bases) and different error rate (i.e., e = 10% and 20%) randomly generated. Regarding the evaluation with real datasets, we use a first set of sequences generated by the Human Pangenome Reference Consortium (Miga and Wang, 2021), consisting of long reads sequenced using Oxford Nanopore Technologies (ONT), PromethION platform, with an average error rate of 5-10%. The sequences are derived from the human cell line HG002, subset to chromosome 12, and restricted to those at least 10 kbp long, for a total number of 1312 sequence pairs of average length equal to 172 kbp (maximum ~306 kbp). Additionally, we use a second dataset comprising ONT MinION reads from Bowden *et al.* (2019), with an average error rate of 5% and restricted to those at least 500 kbp long, for a total number of 48 sequence pairs of average length equal to 630 kbp (maximum ~1 Mbp).

All the executions are performed single-thread on a node running CentOS Linux (release 8.1.1911) equipped with an AMD EPYC 7742 CPU and 1TB of DRAM (distributed in 16 dimms x 64GiB @3200MHz).

### 3.2 Evaluation on simulated data

Table 1 shows the performance results (i.e., execution time and memory) for the different methods using simulated datasets. Overall, the results show that BiWFA is faster and uses less memory than all other methods in the ‘Gap-affine Exact’ category in all cases. In particular, BiWFA requires 713 — 5304× less memory than KSW2-Z2, while being 1.5 — 4.6× faster. Compared to original WFA-based methods (i.e., WFA-high and wfalm), BiWFA uses 478 — 9581 × less memory, being 1.3 — 4× faster. Similarly, BiWFA outperforms the other memory-efficient WFA-based methods (i.e., WFA-med, WFA-low, wfalm-low, and wfalm-rec), requiring 2 — 281× less memory, while being 3.5 — 25.8× faster. In general, most of the pairwise alignment methods evaluated fail to scale megabases-long sequences, requiring more memory than available in the node (i.e., 1TB); as opposed, BiWFA only requires a few hundred MBs of memory.

**Fig. 2:**
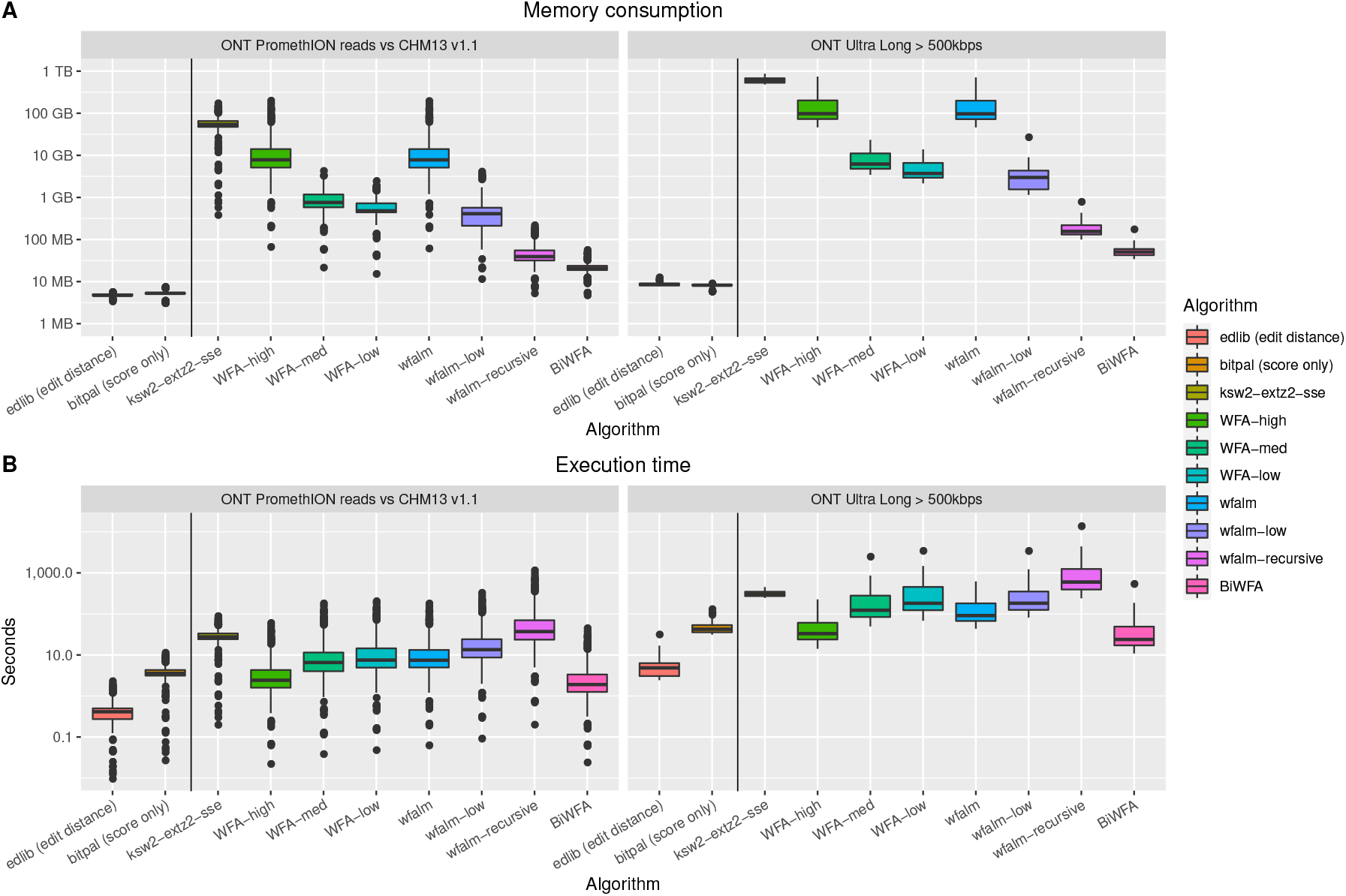
Experimental results from the application of BiWFA and other state-of-the-art libraries and tools on long sequences. Each plot shows the different algorithms versus the memory consumption (A) and execution time (B). We differentiate algorithms that compute edit distance or only the score (left of vertical line) from those that compute the full alignment, CIGAR string included (right of vertical line). In the ONT Ultra Long dataset, not all alignments have been completed, as the tools ran out of memory (OOM) for some of the 48 sequence pairs. In particular, we obtained 10 OOMs with ksw2-extz2-sse, and a single OOM for both WFA-high and wfalm.

**Table 1.** Time and memory performance of pairwise alignment methods on simulated data.

|  | Time (seconds) |  |  |  |  |  |  |  | Memory (MBs) |  |  |  |  |  |  |  |
| --- | --- | --- | --- | --- | --- | --- | --- | --- | --- | --- | --- | --- | --- | --- | --- | --- |
|  | 100 Kbp |  | 500 Kbp |  | 1 Mbp |  | 2 Mbp |  | 100 Kbp |  | 500 Kbp |  | 1 Mbp |  | 2 Mbp |  |
|  | 10% | 20% | 10% | 20% | 10% | 20% | 10% | 20% | 10% | 20% | 10% | 20% | 10% | 20% | 10% | 20% |
| edlib | 0.2 | 0.4 | 4.4 | 9.4 | 17.1 | 33.5 | 67.4 | 132.4 | 4 | 4 | 8 | 8 | 12 | 13 | 22 | 22 |
| bitpal | 1.2 | 1.2 | 31.1 | 31.7 | 123.7 | 123.6 | 499.6 | 493.9 | 3 | 5 | 6 | 8 | 9 | 11 | 14 | 14 |
| ksw2-extz2 | 9.7 | 9.8 | 241.5 | 244.8 | n/a | n/a | n/a | n/a | 19054 | 19084 | 476846 | 477096 | n/a | n/a | n/a | n/a |
| WFA-high | 2.8 | 8.3 | 69.1 | 235.2 | 297.8 | n/a | n/a | n/a | 9005 | 26919 | 229655 | 703138 | 935918 | n/a | n/a | n/a |
| WFA-med | 7.6 | 23.4 | 303.4 | 773.5 | 1935.3 | 3370.3 | n/a | n/a | 827 | 1611 | 12109 | 11521 | 42464 | 24924 | n/a | n/a |
| WFA-low | 9.0 | 27.1 | 544.8 | 1008.5 | 3922.5 | 4959.7 | 15425.2 | 19656.0 | 548 | 878 | 7323 | 5864 | 25409 | 12551 | 52551 | 26067 |
| wfalm | 8.4 | 24.8 | 204.1 | 571.6 | 789.5 | n/a | n/a | n/a | 8994 | 26857 | 225631 | 670617 | 902513 | n/a | n/a | n/a |
| wfalm-low | 15.8 | 47.1 | 382.5 | 1098.5 | 1464.4 | 4210.1 | 5722.6 | 16763.8 | 443 | 822 | 4546 | 8668 | 10471 | 30805 | 36420 | 69245 |
| wfalm-rec | 44.8 | 133.3 | 1345.1 | 4137.7 | 5963.6 | 17356.1 | 23450.9 | 72761.7 | 41 | 74 | 243 | 420 | 500 | 908 | 1100 | 1896 |
| BiWFA | **2.1** | **6.3** | **52.1** | **166.4** | **226.5** | **676.9** | **923.2** | **2846.3** | **19** | **27** | **62** | **90** | **98** | **186** | **202** | **247** |

Execution time (in seconds) and memory (in MBs) required by the different pairwise alignment algorithms on simulated datasets. Executions that failed appear as "n/a". Best performing method in the 'Gap-affine Exact' category are marked in bold. Although Edlib and BitPal are not directly comparable to the other methods, we included them in the comparison as a reference.

### 3.3 Evaluation on real data

Figure 2 shows the performance results obtained for all the evaluated algorithms in terms of execution time and consumed memory. BiWFA uses many times less memory than other methods. In particular, when aligning ultra long ONT sequences (Figure 2B), BiWFA requires between 68 — 93 × less memory compared to wfalm and WFA low-memory modes. Furthermore, BiWFA uses 3.5 × less memory compared to the efficient recursive mode from wfalm (most memory-efficient gap-affine algorithm to date).

At the same time, BiWFA proves to be one of the fastest tools in aligning long sequences. Using ultra long sequences, our method is 23.5 × faster than wfalm’s recursive mode. Moreover, BiWFA’s execution times are similar to those of BitPal (sometimes even faster, 1 – 1.2 × faster on average) computing exact alignments (not just the score) under the gap-affine model.

## 4 Discussion

As long sequencing technologies improve and high-quality sequence assembly decreases in cost, we anticipate that the importance of pairwise alignment algorithm will continue to increase. To keep up with upcoming improvements in sequencing and genomics, pairwise alignment algorithms need to face crucial challenges in reducing execution time and memory consumption. In this work, we have presented the BiWFA, a gap-affine pairwise alignment algorithm that requires *O*(*ns*) time and *O*(*s*) space, being the first algorithm to improve the long standing space lower bound of *O*(*n*). The BiWFA answers the pressing need for sequence alignment methods capable to scaling to genome-scale alignments and full pangenomes.

Most notably, BiWFA execution times are very similar, or even better, than those of the original WFA (despite BiWFA requiring 2958× and 604× less memory when aligning ultra long MinION and PromethION sequences, respectively). This result can be better understood considering the memory inefficiencies that the original WFA experiences when using a large memory footprint. As the sequence’s length and error increases, the original WFA uses a substantially larger memory footprint, putting a significant pressure on the memory hierarchy of the processor. Due to the pervasive memory inefficiencies of modern processors executing memory intensive applications, the original WFA’s performance is severely deteriorated when aligning long sequence datasets (like those from Nanopore presented in the evaluation). In contrast, BiWFA relieves this memory pressure using a minimal memory footprint. As a result, BiWFA is able to balance out the additional work induced by BiWFA’s recursion, delivering a performance on-par with the original WFA.

We have presented the BiWFA using gap-affine penalties. Nevertheless, these very same ideas can be translated directly into other distances like edit, linear gap, or piecewise gap-affine. Moreover, it can be easily extended to semi-global alignment (a.k.a. ends-free, glocal, extension, or overlapped alignment) by modifying the initial conditions and termination criterion. At the same time, the BiWFA retains the strengths of the original WFA: no restrictions on the sequences’ alphabet, preprocessing steps, nor prior estimation of the alignment error.

Due to the simplicity of the WFA’s computational pattern, BiWFA’s core functions can be easily vectorized to fully exploit the capabilities of modern SIMD multicore processors. Our implementation, relies on the automatic vectorization capabilities of modern compilers. As a result, the BiWFA implementation can exploit the SIMD capabilities of any processor supported by modern compilers, without rewriting any part of the source code.

Genomics and bioinformatics methods will continue to rely on sequence alignment as a core and critical component. BiWFA paves the way for the development of faster and more accurate tools that can scale with longer and noisy sequences using a minimal amount of memory. In this way, we expect BiWFA to enable efficient sequence alignment at genome-scale in years to come.

## Acknowledgements

We thank Ragnar Groot Koerkamp for reviewing the manuscript and making useful suggestions.

## Funding

This research was supported by the the European Union Regional Development Fund within the framework of the ERDF Operational Program of Catalonia 2014-2020 with a grant of 50% of total cost eligible under the DRAC project [001-P-001723] and Lenovo-BSC Contract-Framework Contract (2020). It was also supported by the Ministerio de Ciencia e Innovacion MCIN AEI/10.13039/501100011033 under contracts PID2020-113614RB-C21 and PID2019-107255GB-C21, by the Generalitat de Catalunya GenCat-DIUiE (GRR) (contracts 2017-SGR-313, 2017-SGR-1328, and 2017-SGR-1414). M.M. was partially supported by the Spanish Ministry of Economy, Industry and Competitiveness under Ramon y Cajal fellowship number RYC-2016-21104. S.M. was supported by Juan de la Cierva fellowship grant IJC2020-045916-I funded by MCIN/AEI/10.13039/501100011033 and by “European Union NextGenerationEU/PRTR”. B.P and J.E. were supported, in part, by the United States National Institutes of Health (award numbers: R01HG010485, U01HG010961, OT2OD026682, OT3HL142481, and U24HG011853). E.G. was supported by NIH/NIDA U01DA047638 and NSF PPoSS Award #2118709. A.G. acknowledges Dr. Nicole Soranzo’s efforts to establish a pangenome research unit at the Human Technopole in Milan, Italy.

## Notes

### Competing Interest Statement

The authors have declared no competing interest.

### Summary of Updates

Added an extended experimental evaluation and a BiWFA example diagram.

