## Supplementary Material for "Optimal gap-affine alignment in *O*(*s*) space"

### 1 Proof of the correctness lemma

In order to reason about the properties of the WFA dynamic programming structures, it is helpful to invoke certain properties of the Needleman-Wunsch dynamic programming matrices. Accordingly, we will provide the recursions here to introduce the notation.

$$\begin{aligned} D_{i,j} &= \min\{M_{i-1,j} + o + e, D_{i-1,j} + e\} \\ I_{i,j} &= \min\{M_{i,j-1} + o + e, I_{i,j-1} + e\} \\ M_{i,j} &= \min\{I_{i,j}, D_{i,j}, M_{i-1,j-1} + x \cdot \mathbb{I}(q[i-1] \neq t[j-1])\}, \end{aligned} \tag{1}$$

where $\mathbb{I}$ is the indicator function that evaluates to 1 if its argument is true and 0 otherwise. The base case of the recursion is $M_{0,0} = 0$. We also adopt the convention that $D_{0,j} = I_{i,0} = \infty$ for all $i$ and $j$. An optimal alignment can be identified with a *traceback path* through these matrices: a sequence of cells that indicate which of the options from the recursion achieved the minimum score.

Before proving the correctness lemma, we prove two useful properties of the Needleman-Wunsch matrices.

**Lemma 1.** *$M$ is monotonically non-decreasing along each diagonal.*

*Proof.* Choose integers $i$ and $j$ such that $0 \leq i < m$ and $0 \leq j < n$, and we will show that $M_{i,j} \leq M_{i+1,j+1}$, which is sufficient to prove the claim. $M_{i+1,j+1}$ corresponds to the score of an optimal alignment of $q_{0:i}$ and $t_{0:j}$. Any traceback path of this alignment must include a coordinate $(i, y)$ with $y \leq j$ or $(x, j)$ with $x \leq i$. Without loss of generality, assume that there is an optimal alignment path that includes $(i, y)$, and choose $y$ to be the maximal such value within this path. We consider two cases:

1. $y = j$. Then $(i, j)$ is on the traceback path from $M_{i+1,j+1}$ and hence $M_{i,j} \leq M_{i+1,j+1}$.
2. $y < j$. Then there must be at least $j - y$ horizontal transitions on the traceback path following $(i, y)$ for it to end in diagonal $i - j$. Moreover, since $y$ is chosen to be maximal, $(i, y + 1)$ is not on the traceback path, and there must therefore be at least one gap opened after $(i, y)$. This implies $M_{i+1,j+1} \geq M_{i,y} + o + (j - y)e$. We also have $M_{i,j} \leq M_{i,y} + o + (j - y)e$, since it is possible to reach $(i, j)$ by taking $j - y$ horizontal transitions starting from $(i, y)$.

□

**Lemma 2.** *$D$ and $I$ are monotonically non-decreasing along each diagonal, excluding the boundaries $D_{0,\cdot}$ and $I_{\cdot,0}$.*

*Proof.* The proofs for $I$ and $D$ are essentially identical, so we will prove the claim only for $I$. The argument will be proved by induction on decreasing values for the diagonal $k$. The base case $k = m - 1$ is trivially true because there is only one cell in $I$ in this diagonal (excluding the boundary). Consider $i$ and $j$ such that $0 \le i < m$ and $0 < j < n$, and assume that the induction hypothesis holds for all diagonals $k > i - j$. We will show that $I_{i,j} \le I_{i+1,j+1}$, which is sufficient prove the induction claim for $k = i - j$. Consider two cases.

1. $I_{i+1,j+1} = M_{i+1,j} + o + e$. Then, by Lemma 1, we have
$$I_{i,j} \le M_{i,j-1} + o + e \le M_{i+1,j} + o + e = I_{i+1,j+1}. \tag{2}$$
2. $I_{i+1,j+1} = I_{i+1,j} + e$. Then, by the induction hypothesis, we have
$$I_{i,j} \le I_{i,j-1} + e \le I_{i+1,j} + e = I_{i+1,j+1}. \tag{3}$$

□

We are now equipped to prove the central lemma that demonstrates correctness.

**Lemma 2.1 (from main text).** *The optimal alignment score $s_{opt} \le s$ if and only if there exist $s_f$, $s_r$, and $k$ such that $|s_f - s_r| \le p$ and at least one of the following is true:*

1. $s_f + s_r = s$ *and* $\overrightarrow{\mathcal{M}}_{k,s_f} \ge \overleftarrow{\mathcal{M}}_{k,s_r}$
2. $s_f + s_r = s + o$ *and* $\overrightarrow{\mathcal{I}}_{k,s_f} \ge \overleftarrow{\mathcal{I}}_{k,s_r}$
3. $s_f + s_r = s + o$ *and* $\overrightarrow{\mathcal{D}}_{k,s_f} \ge \overleftarrow{\mathcal{D}}_{k,s_r}$,

*and further, $\overleftarrow{\mathcal{M}}_{k,s_r}$ (resp. $\overleftarrow{\mathcal{I}}_{k,s_r}$, $\overleftarrow{\mathcal{D}}_{k,s_r}$) is included in the traceback of an alignment with score at most $s$ if the first (resp. second, third) condition is true.*

*Proof.* ($\Rightarrow$) Let $(i, j)$ be a coordinate along some optimal traceback path where the dynamic programming value has the minimum difference from $s_{opt}/2$. If there are ties, choose the first among the coordinates that achieve the minimum. We consider three exhaustive cases. In each of them, our goal will be to produce the values $s_f$, $s_r$, and $k$ as required by the claim.

1. *The path is in $M$ at $(i, j)$.* Then the path up to $(i, j)$ and the path after $(i, j)$ correspond to partial alignments in the forward and reverse direction respectively, and their scores are $s_f = M_{i,j}$ and $s_r = s_{opt} - M_{i,j}$. Taking $k = i - j$, we know that the f.r. points in the $k$-th diagonal must be at least as far as this coordinate in their respective directions: $\overrightarrow{\mathcal{M}}_{k,s_f} \ge i \ge \overleftarrow{\mathcal{M}}_{k,s_r}$.

   Because adjacent positions in an optimal traceback path can differ by at most $p$, we have both $|s_f - s_{opt}/2| \le p/2$ and $|s_r - s_{opt}/2| \le p/2$. These imply $|s_f - s_r| \le p$ by the triangle inequality.
2. *The path is in $I$ at $(i, j)$ and not also in $M$ at $(i, j)$.* Then $(i, j)$ is part of a gap that begins at $(i, j')$ for some $j' < j$ and ends at $(i, j' + \ell)$ where $j' + \ell > j$, else the path is also in $M$ at $(i, j)$. Consider the quantity $x = (s_{opt} - 2M_{i,j'})/2e$ across three cases.

   2.1. $x \le 1/2$. Let $s_f = M_{i,j'}$ and $s_r = s_{opt} - M_{i,j'} + o$. These correspond to the scores of the partial alignments before and after $(i, j')$, respectively. Therefore we take $k = i - j'$, and, as previously, the f.r. points within this diagonal must obey the inequality $\overrightarrow{\mathcal{M}}_{k,s_f} \ge i \ge \overleftarrow{\mathcal{M}}_{k,s_r}$.
   Note that $M_{i,j'} \le s_{opt}/2$ else $I_{i,j'}$ would not achieve the minimum difference from $s_{opt}/2$. This implies $x \ge 0$, and in particular $|x| \le 1/2$. Therefore,
   $$|s_f - s_r| = |s_{opt} - 2M_{i,j'} + o| \le |o + 2ex| \le o + 2e|x| \le o + e \le p. \tag{4}$$

2.2. $1/2 < x < \ell - 1/2$. Let $x^*$ be the nearest integer to $x$, and let $s_f = I_{i,j'+x^*}$ and $s_r = s_{opt} - I_{i,j'+x^*} + o$. These correspond to the scores of the partial alignments before and after $(i, j' + x^*)$, respectively. Therefore we take $k = i - j' - x^*$, and, as previously, the f.r. points within this diagonal must obey the inequality $\overrightarrow{\mathcal{I}}_{k,s_f} \geq i \geq \overleftarrow{\mathcal{I}}_{k,s_r}$.
Noting that $|x - x^*| \leq 1/2$ by construction, we also have

$$|s_f - s_r| = |s_{opt} - 2M_{i,j'} - 2x^*e| \leq 2e|x - x^*| \leq e \leq p. \tag{5}$$

2.3. $x \geq \ell - 1/2$. Let $s_f = I_{i,j'+\ell}$ and $s_r = s_{opt} - I_{i,j'+\ell} + o$. These correspond to the scores of the partial alignments before and after $(i, j' + \ell)$, respectively. Therefore we take $k = i - j' - \ell$, and, as previously, the f.r. points within this diagonal must obey the inequality $\overrightarrow{\mathcal{I}}_{k,s_f} \geq i \geq \overleftarrow{\mathcal{I}}_{k,s_r}$.
Noting that $s_{opt}/2 \leq I_{i,j'+\ell}$ else $j \geq j' + \ell$, and also that $I_{i,j'+\ell} = M_{i,j'} + o + \ell e$, we can obtain

$$\begin{aligned} s_{opt} &\leq 2M_{i,j'} + 2o + 2\ell e \\ s_{opt} - M_{i,j'} - \ell e &\leq M_{i,j'} + 2o + \ell e \\ s_r &\leq s_f + o. \end{aligned} \tag{6}$$

Since $x \geq \ell - 1/2$, we also have

$$\begin{aligned} s_{opt} - 2M_{i,j'} &\geq (2\ell - 1)e \\ s_{opt} - M_{i,j'} - \ell e &\geq M_{i,j'} + (\ell - 1)e \\ s_r &\geq s_f - o - e. \end{aligned} \tag{7}$$

These together imply $|s_f - s_r| \leq o + e \leq p$.

3. *The path is in $D$ at $(i, j)$ and not also in $M$ at $(i, j)$.* Same as the previous case.

($\Leftarrow$) We consider the three conditions separately.

1. Let $(i_1, j_1)$ be the coordinates in $M$ corresponding to $\overrightarrow{\mathcal{M}}_{k,s_f}$ and likewise $(i_2, j_2)$ for $\overleftarrow{\mathcal{M}}_{k,s_r}$. The partial alignments corresponding $M_{i_2,j_2}$ and $\overleftarrow{\mathcal{M}}_{k,s_r}$ can be concatenated into a full alignment with score $M_{i_2,j_2} + s_r$. By Lemma 1, this score is at most $M_{i_1,j_1} + s_r = s_f + s_r = s$.

2. Let $(i_1, j_1)$ be the coordinates in $I$ corresponding to $\overrightarrow{\mathcal{I}}_{k,s_f}$ and likewise $(i_2, j_2)$ for $\overleftarrow{\mathcal{I}}_{k,s_r}$. The partial alignments corresponding $I_{i_2,j_2}$ and $\overleftarrow{\mathcal{I}}_{k,s_r}$ can be concatenated into a full alignment with score $I_{i_2,j_2} + s_r - o$. By Lemma 2, this score is at most $I_{i_1,j_1} + s_r - o = s_f + s_r - o = s$.

3. Same as previous condition.

□
